# Generating hard-to-obtain information from easy-to-obtain information: applications in drug discovery and clinical inference

**DOI:** 10.1101/2020.08.20.259598

**Authors:** Matthew Amodio, Dennis Shung, Daniel Burkhardt, Patrick Wong, Michael Simonov, Yu Yamamoto, David van Dijk, Francis Perry Wilson, Akiko Iwasaki, Smita Krishnaswamy

## Abstract

In many important contexts involving measurements of biological entities, there are distinct categories of information: some information is easy-to-obtain information (EI) and can be gathered on virtually every subject of interest, while other information is hard-to-obtain information (HI) and can only be gathered on some of the biological samples. For example, in the context of drug discovery, measurements like the chemical structure of a drug are EI, while measurements of the transcriptome of a cell population perturbed with the drug is HI. In the clinical context, basic health monitoring is EI because it is already being captured as part of other processes, while cellular measurements like flow cytometry or even ultimate patient outcome are HI. We propose building a model to make probabilistic predictions of HI from EI on the samples that have both kinds of measurements, which will allow us to generalize and predict the HI on a large set of samples from just the EI. To accomplish this, we present a conditional Generative Adversarial Network (cGAN) framework we call the Feature Mapping GAN (FMGAN). By using the EI as conditions to map to the HI, we demonstrate that FMGAN can accurately predict the HI, with heterogeneity in cases of distributions of HI from EI. We show that FMGAN is flexible in that it can learn rich and complex mappings from EI to HI, and can take into account manifold structure in the EI space where available. We demonstrate this in a variety of contexts including generating RNA sequencing results on cell lines subjected to drug perturbations using drug chemical structure, and generating clinical outcomes from patient lab measurements. Most notably, we are able to generate synthetic flow cytometry data from clinical variables on a cohort of COVID-19 patients—effectively describing their immune response in great detail, and showcasing the power of generating expensive FACS data from ubiquitously available patient monitoring data.

**Bigger Picture:** Many experiments face a trade-off between gathering easy-to-collect information on many samples or hard-to-collect information on a smaller number of small due to costs in terms of both money and time. We demonstrate that a mapping between the easy-to-collect and hard-to-collect information can be trained as a conditional GAN from a subset of samples with both measured. With our conditional GAN model known as Feature-Mapping GAN (FMGAN), the results of expensive experiments can be predicted, saving on the costs of actually performing the experiment. This can have major impact in many settinsg. We study two example settings. First, in the field of pharmaceutical drug discovery early phase pharmaceutical experiments require casting a wide net to find a few potential leads to follow. In the long term, development pipelines can be re-designed to specifically utilize FMGAN in an optimal way to accelerate the process of drug discovery. FMGAN can also have a major impact in clinical setting, where routinely measured variables like blood pressure or heart rate can be used to predict important health outcomes and therefore deciding the best course of treatment.

## 1. Introduction

When collecting information on biological entities, for example hospital patients, cells, or drugs, we are often faced with the choice of collecting easy-to-obtain information (EI) on many entities or collecting hard-to-obtain information (HI) on a few entities. For example, in a drug library of millions of drugs, it is easy to obtain chemical structure information but hard to obtain RNA sequencing information of cells treated with drugs. On patients, it may be easy to obtain information such as heart rate and lab values, but hard to obtain blood flow cytometry information. Here, we present a neural network-based method that can bridge the gap between these sources of information on entities like drugs or patients.

We introduce a framework based on a conditional Generative Adversarial Network (cGAN) that we call Feature Mapping GAN (FMGAN), which learns a mapping from EI to a distribution of HI. The FMGAN takes in noise as input, the EI information as the *condition* and the HI as the output. For instance, given the chemical structure of a drug, we can build a mapping to the RNA sequencing of cells under the drug. Here, the EI is the chemical structure and is used as the condition for the cGAN. Corresponding HI is then produced by the generator of the cGAN. We showcase this in many settings involving different information obtained on drugs and patients.

Our use of a GAN-based framework is motivated by our applications’ having complex, one-to-many relationships between the EI and the HI. To illustrate this further, consider a simple linear mapping between an EI variable and an HI variable. The linearity guarantees that small changes in the EI will result in a small change in the HI, i.e. the mapping is *smooth*. However, with chemical structure, for example, this is known not to be true: a small change in chemical structure can lead to vastly different properties of a drug. Non-linear mappings can also be simple, such as a simple threshold decision: if a particular clinical variable completely determines patient outcome, a logical decision with a threshold would suffice. However, clinical outcomes are the result of complex couplings between large groups of variables. This necessitates a rich mechanism of mapping EI to HI, capable of representing the necessary complexity. Moreover, the mapping has to be stochastic. Since, it is unlikely that the EI has complete information about the drug or patient in question, it is important for each EI condition to be able to map to a range or a distribution of HI conditions. For example, replicates of a drug perturbation experiment result in different gene expression results even when applied on the same cell line [1]. This stochastic response can only be captured by a generative model that can produce stochastic output. As GANs learn complex mappings from a random noise space (and, for cGANs, an EI space) to the HI space, they have the required complexity and stochasticity. And with their flexible training paradigm, they do so without having to make strong assumptions like those involved in choosing a parametric family for the form of the HI distribution.

One of our motivating examples through this paper is the **drug discovery process**. A major part of pharmacological research is devoted to drug discovery, where a large number of drug compounds have to be sorted to find a small number of promising candidates [2]. This search can be guided by information about the drug itself, as well as by the past history of how other drugs have performed [3]. By looking for drugs similar to ones that have shown success previously, promising candidates with improved toxicity or efficacy can be identified. Improvements in this form of research, called hit-to-lead, can save significant time and money. The search for promising candidate drugs is a daunting task, since the state space of molecular libraries is in the millions, and possible drugs is in the tens of thousands or more [4].

Here we specifically consider measurements involving drug perturbations, a commonly used technique for measuring the effect of a drug [5–7]. We utilize drug perturbation data from the L1000 Connectivity Map dataset [1]. Perturbation involves introducing the drug to a sample of cells and then measuring the gene expression of those cells after the drug treatment. By comparing the gene expression of the cells before and after drug treatment, researchers can infer information about what the drug does and how it works. Because perturbing a cell line with a drug involves physically performing an experiment, including obtaining the cells, applying the drug, and getting the sequencing results, this process can be expensive and time-consuming. We use the FMGAN to generate the RNA-sequencing results from the drug structure to speed this process up by not having to perform all of the experiments exhaustively. If only a subset of the drugs have a priori RNA-sequencing measurements, the rest can be generated with the FMGAN, obviating the need for additional experimentation on a large number of candidates.

Another motivating setting is that of **clinical data**. In the clinical setting, some measurements are readily available EI, either because they are already measured as part of the standard patient monitoring, or because they are non-invasive and do not pose any risk. We work with two clinical datasets of this type. The first is an Electronic Intensive Care Unit (eICU) Collaborative Research Database dataset, which includes as EI standard clinical measurements such as body temperature, heart monitoring, and standard blood work [8,9]. With this EI we generated predicted clinical mortality, a measurement whose value can normally only be obtained too late to act upon. Rather than due to financial expense, this measurement is hard-to-obtain because it is irreversible, involving patient mortality. With the FMGAN, predictions can be accurately generated from the EI and thus preventative measures can be taken while positive interventions are still possible.

The second clinical dataset we work with uses similar clinical measurements as EI, but this time on COVID-19 patients from Yale New Haven Hospital. In this case, the HI information are future single-cell flow cytometry measurements from samples gathered on some of the patients. In practice, these types of single-cell measurements cannot be performed exhaustively on every patient in the clinic, for reasons of cost as well as time sensitivity. Thus, we use the FMGAN to be able to generate future flow cytometry data which depicts compartments of the immune system from readily available clinical data. With the FMGAN, we are then able to generate flow cytometry data for any number of patients who only have clinical measurements available. This can be valuable as immune responses have been shown to be highly predictive of mortality in COVID-19 [10].

In each of these datasets we not only utilize the natural flexibility of the cGAN in mapping, but also explicitly design mechanisms for the cGAN to take advantage of any structure that *does exist* in the EI. While EI-HI mapping is rarely linear or simple, there are many instances in which the HI is smooth and respects geometric or manifold structure in the EI—which can be explicitly represented. Here, we show two ways of of taking into account latent structure in the EI. The first is by embedding the EI into lower dimensional manifold-intrinsic coordinates, such as with the PHATE dimensionality-reduction method, which has been shown to preserve manifold affinity [11]. We show this on the case of drug perturbations where we measure some genes on perturbed cell lines, and impute the other genes. Since the underlying cellular manifold measured is the same, both measured and withheld genes should respect this structure. We also show this on clinical data where ICU measurements are embedded with PHATE and then the embeddings are used to impute clinical outcomes. The second situation, rather than embedding EI with PHATE, is to use a convolutional neural network to find a latent space embedding of the data. We use this encoding of the EI where it is the chemical structure of the drug. Here, we create a rich set of convolutional features of the chemical structure by treating it as an image. In particular a small change in the structure can be reflected as larger changes in convolutional filter outputs, and thus the latent space has more regularity with respect to the mapping than the original chemical structure space.

## 2. Results

### 2.1. FMGAN

The FMGAN we propose uses a conditional Generative Adversarial Network (cGAN) to generate hard-to-collect information (like sequencing results from a perturbation experiment) from other easy-to-collect information (like basic information on the drugs used). Specifically, we propose a cGAN with the easy-to-collect information as conditions and the hard-to-collect information as the data distribution. A cGAN is a generative model that learns to generate points based on a conditional label that is given to the generator *G*. In the adversarial learning framework, *G* is guided into generating realistic data during training by another network, the discriminator *D*, that tries to distinguish between samples from the real data and samples from the generated data. The generator *G* and discriminator *D* are trained by alternating optimization of G and D.

A standard GAN learns to map from random stochastic input *z* ∼ *N*(0, 1) (or a similarly simple distribution) to the data distribution by training *G* and *D* in alternating gradient descent with the following objective:

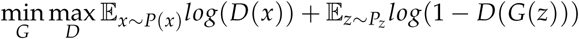

The generator in a cGAN receives both the random stochastic input *z* and a conditional label *l* and thus has the following objective:

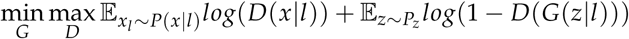

The cGAN was originally used in image generation contexts, where the condition referred to what type of image should be generated (e.g. a dog). The cGAN is useful in this context because the generator *G* receives a sample from a noise distribution (as in a typical GAN) as well as the condition. Thus, it is able to generate a *distribution* that is conditioned on the label, as opposed to a single deterministic output conditioned on the label. In the original use case, it can learn to generate a wide variety of images of dogs when given the conditional label for dogs, for example. While many previous methods exist for generating a single output from a single input, there are few alternatives for generating a distribution of outputs from a single input without placing assumptions on the parametric form of the output distribution.

The framework of the FMGAN is summarized in Figure 1. The columns of the data are separated into easy-to-collect information (EI) and hard-to-collect information (HI). In the notation of the GAN, we use the EI as the conditional label *l* and the HI as the data *x*. For observations that have both, we train the FMGAN with the generator receiving a label *l* and a noise point *z*, while the discriminator receives the label *l* and both real points *x* and the generated points *G*(*z*|*l*). Then, after training, the generator can generate points for conditions *l* without known data *x*. This allows us to impute HI where we only have EI.

**Figure 1.**
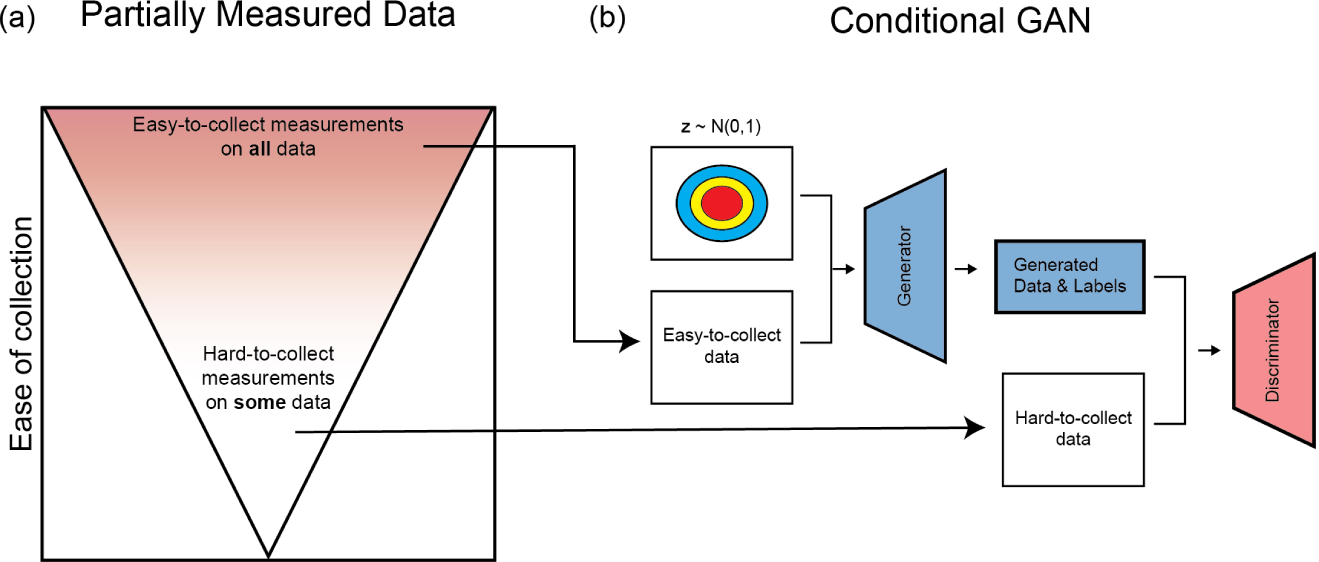
(a) The measurements on data are separated into “easy-to-collect information” (EI) and “hard-to-collect information” (HI). The easy-to-collect measurements are available on all data, while the hard-to-collect measurements are only available on some data. (b) With a Conditional GAN, we can learn to model the relationship between these two categories of measurements.

The FMGAN architecture is designed to take advantage of complex relationships between the condition space and the data space. A single underlying entity (e.g. a drug or a patient) has a representation in both spaces. In the EI space, the drug is a point, while in the HI space the drug is represented by a distribution of cells perturbed by it. Despite the difference in structure, the FMGAN is able to leverage regularities in the relationship between the two spaces. This relies on the FMGAN being able to leverage manifold structure inherent within each space (for more discussion of manifold structure, please see the supplementary information).

In some cases, the data modality for the EI is difficult to utilize: for example, the chemical structure of the drug. The chemical structure can be represented as a string sequence called SMILES or a two-dimensional image of the structure diagram. Small changes in the chemical structure can have large changes on its function, but may appear to be minor changes to the overall SMILES string or the overall structure diagram image. Thus, we use an embedding neural network, parameterized as a convolutional network, to process these representations into a more regular space where standard distances and directions are meaningful. This parameterization is crucial, as originally the structure is not linear (or else simpler models could leverage it). But with convolutional networks, small changes in the input can cascade down into deeper layers in complex ways and make potentially large, meaningful shifts in the embedding. We further detail the architecture and design of this network in the Methods section.

### 2.2. Modeling drug perturbation experiments

We first demonstrate the results of our FMGAN model on data from the L1000 Connectivity Map (CMap) dataset [1]. The CMap dataset contains a matrix of genes by count values on various cell lines under different drug perturbations. We examine the A375 cell line, a cell line from a human diagnosed with malignant melanoma. In this densely measured dataset, we have all gene expression measurements for each drug. Each drug also has various numbers of replicates of the same experiment. These replicates produce variable effects, motivating the need for a framework that is capable of modeling such stochasticity.

We design four separate experiments with this dataset:

1. A proof-of-concept that the cGAN framework can effectively model and predict gene expression values when the conditions are known to be meaningful because they are selected holdout genes from the expression matrix itself.
2. An experiment where the conditions are taken from a non-linear dimensionality reduction method applied to the expressions.
3. A test of the full FMGAN pipeline where conditions represent chemical structure in the form of SMILES strings, and thus embeddings for conditions must be learned.
4. A variation of the chemical structure conditions where they are represented as images of structure diagram.

In each dataset, the measurement we choose for evaluation is maximum mean discrepancy (MMD) [12]. We choose this because we require a metric that is a distance between distributions, not a distance merely between points. Taking the mean of a distance between points would not capture the accuracy of any moments in the desired distribution beyond the first one. For the experiments based on drug metadata (the SMILES strings and the chemical structure images experiments), we consider the drug’s distribution to be all of the gene profiles from that drug. For the experiments with conditions derived from each gene profile (the heldout genes and dimensionality-reduction experiments), we take a neighborhood of drugs around each condition and compare the predicted distribution of gene profiles for those drugs with the true distribution.

We compare our FMGAN to a baseline not built off of the cGAN framework. In developing a baseline, we must compare to a model that takes in a point and outputs an entire distribution. As most existing work yields deterministic output, we create our own stochastic distribution yielding model to compare to. This model, which we term simply “Baseline”, takes a condition and a sample from a random noise distribution as input, just like our FMGAN. However, unlike our model which uses adversarial training and a deep neural network, the Baseline is a simpler, feed forward neural network that minimizes the mean-squared-error (MSE) between the output of a linear transformation and the real gene profile for that condition. As it is given noise input as well as a condition, it is still able to generate whole distributions as predictions for each condition, rather than deterministic single points. As generating conditional distributions (especially based off of oddly structured conditions like images or strings) is relatively understudied in the computational biology field, we find no directly comparably published methods that can be applied to this problem, thus necessitating our creating Baseline.

#### 2.2.1. Predicting gene expression under drug perturbation

To show our cGAN can learn informative mappings from the EI space to the gene expression space, as distinct from the rest of the process, we first choose a means of obtaining EI that are known to be meaningfully connected to the gene expression space. Specifically, we artificially hold out ten genes and use their values as EI, with the GAN tasked with generating the values for all other genes. This experimental design is summarized in Figure 2a. We choose the ten genes algorithmically by selecting one randomly and then greedily adding to the set the one with the least shared correlation with the others, to ensure the information in their values have as little redundancy as possible: PHGDH, PRCP, CIAPIN1, GNAI1, PLSCR1, SOX4, MAP2K5, BAD, SPP1, and TIAM1. In addition to dividing up the gene space to use these ten genes to predict all of the rest, we also divide up the cell space and train on 80% of the cell data, with the last 20% heldout for testing.

**Figure 2.**
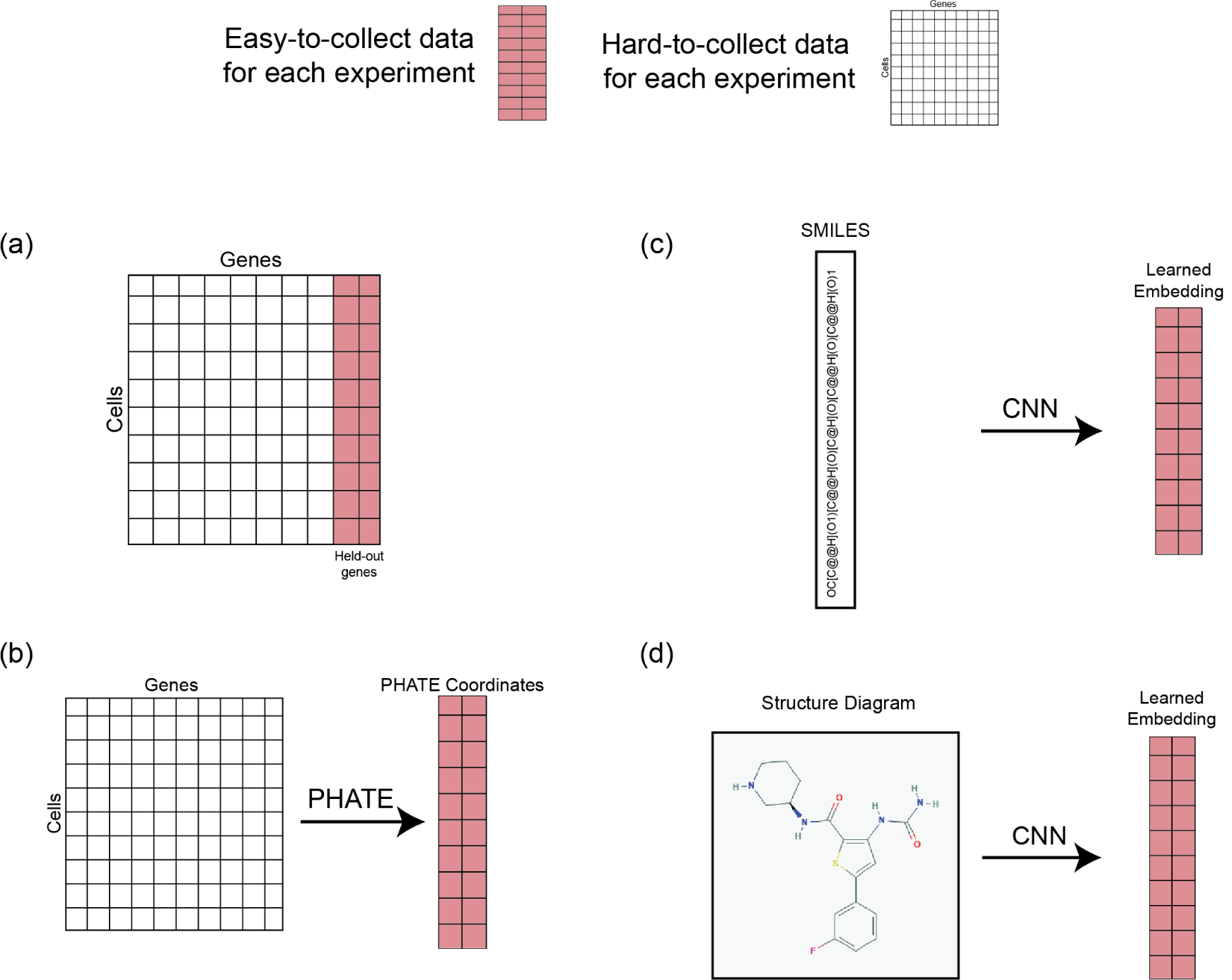
The formation of easy-to-collect (red columns) and hard-to-collect (white columns) data for each experiment with drug perturbation data. (a) in the held-out genes experiment, the easy-to-collect measurements are taken from held-out genes (b) in the PHATE coordinate experiment, they are the result of running on the genes matrix (c) in the SMILES string experiment, the easy-to-collect data is an embedding from processing this representation with a CNN (d) in the structure diagram experiment, it is the same as in the SMILES string experiment except run on the structure diagrams.

We find our cGAN is able to successfully leverage information in the EI space to accurately model the data. We designed our proof of concept deliberately so that the true values are known for each gene expression and drug we ask our network to predict. These values can be compared to the predictions with MMD for a measure of accuracy.

Our cGAN is able to generate predictions with an MMD of 2.847 between it and the validation set (drugs it has never previously seen), showing it very effectively learned to model the dependency structure between the EI space and the HI space, even on newly introduced drugs (Table 1). This is in comparison to the Baseline model, which has a higher (worse) MMD of 2.922. It is noteworthy that the FMGAN outperforms the baseline even in this case, where no processing of the EI needs to take place, as they are numerically meaningful values to begin with.

**Table 1.** MMD scores (lower is better) across all datasets for the drug data for both models. The FMGAN more accurately predicts the distribution from each condition for all methods of forming the condition space.

| MMD Scores | Heldout Genes | PHATE Coordinates | SMILES | Chemical Structure Image |
| --- | --- | --- | --- | --- |
| FMGAN | **2.847** | **0.179** | **1.191** | **1.177** |
| Baseline | 2.922 | 0.330 | 1.510 | 1.798 |

We also can visualize the embedding spaces learned by the generator to investigate the model. Shown in Figure 3a are the generator’s embeddings colored by each of the heldout genes. As we can see, the generator found some of these more informative in learning an EI embedding than others. We can quantify this by building a regression model to try to predict the value of each gene given the embedding to determine the most valuable of the heldout genes. By this measure, PHGDH, PRCP, and GNAI1 are the most important genes. Analyzing the embeddings in this way is useful for determining which part of the EI space was most informative for generating the HI space, and we will continue to do this with more complex EI in later experiments.

**Figure 3.**
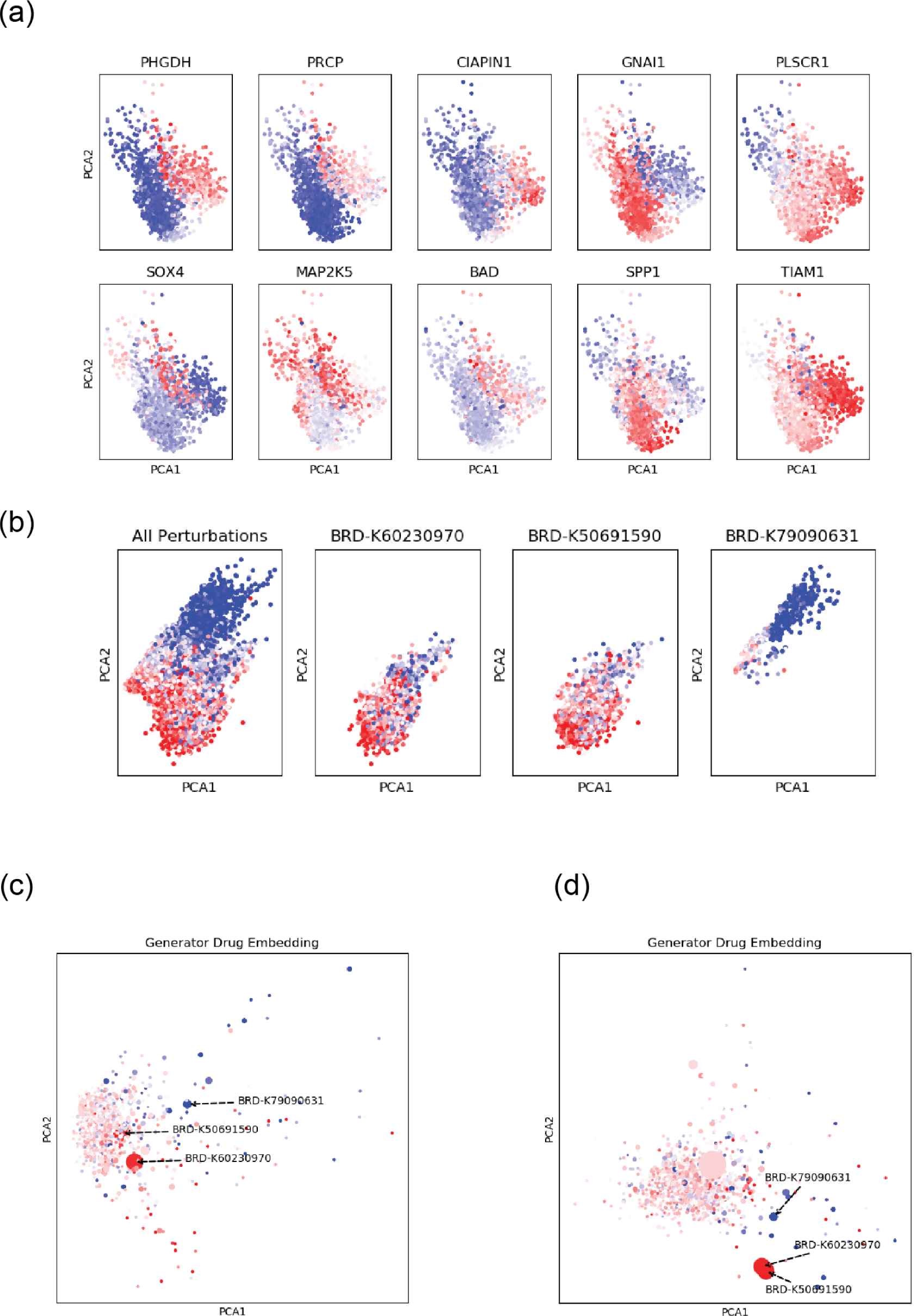
(a) Visualization of the embedding of *cells* in the held-out genes experiment, colored by each held-out gene. The network has inferred the structure of the space from these genes. (b) The raw data, colored by the expression of gene EIF4G2, separated into the three most abundant drugs: BRD-K60230970, BRD-K50691590, and BRD-K79090631. (c) The generator’s embedding space of *drugs* from the SMILES strings experiment, with the same three drugs highlighted. The embedding in shows that the drugs with similar distributions have been embedded into similar locations in the learned embedding space. (d) The same as in (c) but with the structure diagram experiment.

#### 2.2.2. PHATE coordinates as conditions for manifold-structured EI

Our next experiment formulates the EI space not as individual heldout genes, but instead on a dimensionality-reduced representation of the whole space. We theorize that this approach would be beneficial over the previous held-out-genes experiment if the EI data exhibits manifold structure. If it does, this processing will have made a geometric representation of the EI that corresponds to the HI, and thus the mapping is computationally simpler. Previous work has shown that gene expression profiles often do exhibit this manifold structure [11,13,14].

We run the embedding tool PHATE on the gene profiles to calculate two coordinates, which we then use as EI in our FMGAN [11]. Doing so preserves the manifold structure of the data, allowing for a meaningful transformation to the HI space. This process is depicted in Figure 2b. As usual, we separate cells into an 80%/20% training/testing split for evaluation purposes, after being subsampled to ten thousand points for computational feasibility with the dimensionality reduction method, and we report scores on the evaluation points.

As shown in Table 1, once again the FMGAN better models the target distribution, as measured by MMD between its predictions in the neighborhood of each point and the true values. The FMGAN’s predictions had an MMD of 0.179, compared to the baseline MMD of 0.330 (a 45.7% improvement). It is also interesting to note that although the MMDs are not directly comparable across the experiments (because the target distribution is changing each time, from all drugs in neighborhood around a coordinate to all drugs with the same metadata), the PHATE coordinates provide the most accurate predictions.

#### 2.2.3. Predicting Gene Expression from Drug Chemical Structure

Next, we test the full pipeline of FMGAN by using SMILES string embeddings as the EI (summarized in Figure 2c). This is a much more challenging test case, because in the previous cases each point in HI space had a distinct condition, and in the case of the PHATE coordinates, that condition was derived from the data it had to predict. In this case, many different data points have the same condition, and thus the relationship is much less direct between the EI and the HI.

An additional wrinkle also arises in this setting where the conditions to the cGAN are learned from a raw data structure, rather than *a priori* existing in their final numerical form like heldout genes or PHATE coordinates. Since *G* and *D* are trained adversarially and each depends on the embedder *E*, the networks could try to beat each other by manipulating the embeddings into being non-informative for the other network. Thus, we let *G* and *D* learn their own embedder *E*, thus removing the incentive to make *E* non-informative.

As in the previous experiment, we separate the data into an 80%/20% training/testing split for evaluation purposes, but this time split along the drugs since each condition gives rise to many points in the HI space. Table 1 indicates that the FMGAN had an MMD of 1.191 compared to the baseline of 1.510 (a 21.1% improvement).

##### EI Space Analysis

In this section, we investigate further the EI space learned from the SMILES strings by the generator. In the two previous experiments, the conditions given to the FMGAN had information more readily available, either in the form of raw data or even more informative PHATE coordinates. The SMILES strings, by contrast, must be informatively processed for the learned conditions to be meaningful.

In this learned EI space, there is one condition coordinate for each drug (while the HI consists of many perturbations from each drug). Shown in Figure 3b is the raw data colored by the value of gene EIF4G2. Then, all of the perturbations from each of three drugs are shown separately: BRD-K60230970, BRD-K50691590, and BRD-K79090631. As we can see, the first two are characterized by high expression of this gene and are quite similar to each other. The third, however, is quite distinct, in a separate space of the embedding, and is characterized by much lower expression of this gene.

We compare this to the embedding learned by the generator, which we show in Figure 3c. In this plot, each drug is one point, colored by the mean gene value of all perturbations for that drug and with a point whose size is scaled by the number of perturbations for that drug. We see that the first two drugs are in the central part of the space, and closer to each other than they are to BRD-K79090631. The drug BRD-K79090631 is off in a different part of the space, along with other drugs low in EIF4G2. This shows that the learned conditions from the generator have indeed identified information about the drugs and taken complex sequential representations and mapped them into a much simpler space.

#### 2.2.4. Predicting gene expression from drug structure diagrams

The final experiment we consider for the drug perturbation data is the formation of the condition space from an image representation of the chemical structure (Figure 3d). These images are downloaded from the PubChem PUG REST API [15]. An example image for the drug BRD-U86686840 is shown in Figure 2d. They are given as input to a two-dimensional CNN designed for image processing, as points in the original *h* x *w* x *c* pixel space, with *h* = *w* = 64 and *c* = 3. While a CNN is used in both the SMILES string case and this one, the underlying data is in a fundamentally different structure. As in the SMILES string experiment, both the generator and the discriminator learned their own CNN to develop embeddings adversarially.

Table 1 shows that the FMGAN performed slightly better with these chemical structure diagrams as compared to the SMILES strings (1.177 MMD). The baseline model scored significantly worse with these images as compared to the SMILES strings. This illustrates the FMGAN’s flexibility, as it performs comparably with drastically different structures (a long one-dimensional string as opposed to a natural image). That the chemical structure images perform slightly better is perhaps a sign that two-dimensional image convolutional networks are currently more effective at distilling this information than one-dimensional sequence convolutional networks, but the FMGAN’s flexible framework allows it to keep improving with advances in deep learning architectures. Another possibility is that the structure diagrams have relevant information more easily separable from irrelevant information, making them an easier statistical task.

##### EI Space Analysis

In Figure 3d, we show the learned embedding from the generator. We color the embedding by the same gene and highlight the same three drugs as in the previous experiment: BRD-K60230970, BRD-K50691590, and BRD-K79090631. As before the learned conditions have taken a space where it is hard to characterize the information it contains (raw images in pixel space) and mapped them to a simpler space with numerically meaningful points. This can be seen by noting that the two drugs with similar distributions in the raw data (BRD-K60230970 and BRD-K50691590) have been mapped to nearly identical conditions, while they are separate from the drug with a very different distribution (BRD-K79090631). In fact, this goes towards an explanation of the improvement in performance over the SMILES string model, as the embedder has placed the drugs with similar distributions closer to each other in conditions, making the generator’s job easier.

### 2.3. Predicting clinical outcomes

We demonstrate the versatility of our proposed method by experimenting on data in a very different context from the drug perturbations of the previous section. Here we work on clinical data from two different datasets. In each case, we use data derived from clinical measurements on patients to predict their clinical outcomes.

#### 2.3.1. Predicting eICU clinical outcomes

For our first clinical experiment, we use data from patients at high risk for mortality due to severe illness, selected from the eICU Collaborative Research Database [8,9]. As conditions for the FMGAN, we use measurements that are components of the in the widely-used APACHE score. The APACHE score predicts mortality from age, immunocompromised status, heart measures, and respiratory measures [16]. We pass these features through PHATE to develop conditions and then predict mortality as our response variable. For more details on the data and pre-processing, please see the Supplementary Information.

Figure 4 shows the real data, which is noisy but still shows different density of mortality in different parts of the space. We also see the FMGAN generated data next to it: qualitatively, these predictions resemble the raw data to a substantial degree. As a baseline, we can build a linear regression model that tries to predict this response variable as a function of the coordinates. Due to the probabilistic nature of the response, the linear regression predicts a low chance of mortality everywhere in the space, with a slight uptick in probability in the dense region.

**Figure 4.**
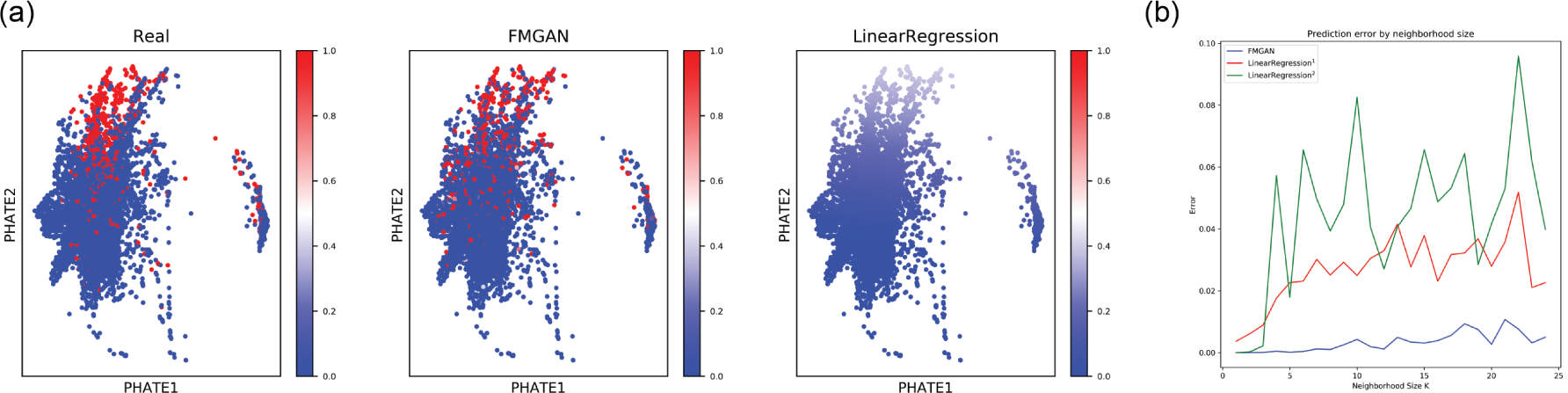
(a) The raw data from the eICU clinical outcomes experiment, along with FMGAN generated data and a linear regression baseline. (b) Quantitative evaluation of the model and the baseline.

This is different from our FMGAN, which better models the binary nature of the output: in each region there are some zeros and some ones as opposed to every point having a small constant value like 0.1. To quantify the accuracy of each model, we have to develop an evaluation criterion that looks at different regions and compares the true number of mortalities and predicted number in that region. This metric assumes that within each local neighborhood, which point gets which label is partially determined by randomness, and that the true signal is the proportion of points within that neighborhood. Using this metric, we can compute the prediction error as the difference between the predicted number of mortalities in a neighborhood and the true number.

Specifically, we compute K partitions of the data using the nearest neighbors clustering algorithm. In each neighborhood, we compare the proportion of positive predictions (using a threshold of 0.5) and the proportion of real positive outcomes with a mean-squared error measurement. Figure 4b shows this for varying numbers of neighborhoods K. Also, while our model naturally outputs data like the underlying data and thus has an easily identifiable threshold of 0.5 for a positive prediction, the linear regression does not have an obvious choice for a threshold for a positive prediction. We use both the default 0.5 (labeled LinearRegression^1^) and the *i*^*th*^ percentile of the output, where *i* is chosen to match the total proportion of real responses equal to one and the predicted proportion responses equal to one.

Figure 4b shows a chart of these values for increasing numbers of neighborhoods to divide the space into. The errors for the linear regression models range from 0.02 to 0.10 depending on the neighborhood size, while the FMGAN remains below 0.01 for all neighborhood sizes. This means the regression model has at least doubled the error of the FMGAN in each neighborhood size. The stochasticity in the data makes it so that the GAN framework, which incorporates stochastic noise input, is best able to generate output like the real data.

#### 2.3.2. Predicting COVID-19 clinical outcomes

In this section we present an experiment that learns a mapping between clinical measurements and FACS measurements from COVID-19 patients [17]. The clinical measurements are taken from the first 24 hours in the ICU, with a patient’s record being the most extreme value taken during that period when more than one record is taken. To test the ability of FMGAN to make practical, and actionable predictions we learn to generate the first flow cytometry measurement, taken from anywhere from the first week to the eleventh week of the stay. Thus, we model *future* flow cytometry with *present* clinical data.

The conditions we use for the FMGAN, as in the previous experiment, are PHATE coordinates of embedded clinical variables. In the PHATE embedding each patient is represented by a a vector of variables, listed in the supplement in Table 3. For each of 129 patients, we also have matched FACS measurements on 14 proteins obtained from each patient, which are listed in the supplement in Table 4. While the clinical measurements are relatively easy and inexpensive to obtain, FACS samples are comparatively expensive and time-consuming to obtain. Thus, we wish to learn a model that can accurately generate FACS data from a patient’s clinical measurements alone.

**Table 2.**
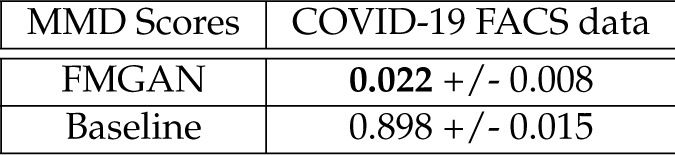
MMD scores (lower is better) on the COVID-19 data, with mean and standard deviation across the 13 held-out patients. The FMGAN outperforms the baseline significantly.

**Table 3.**
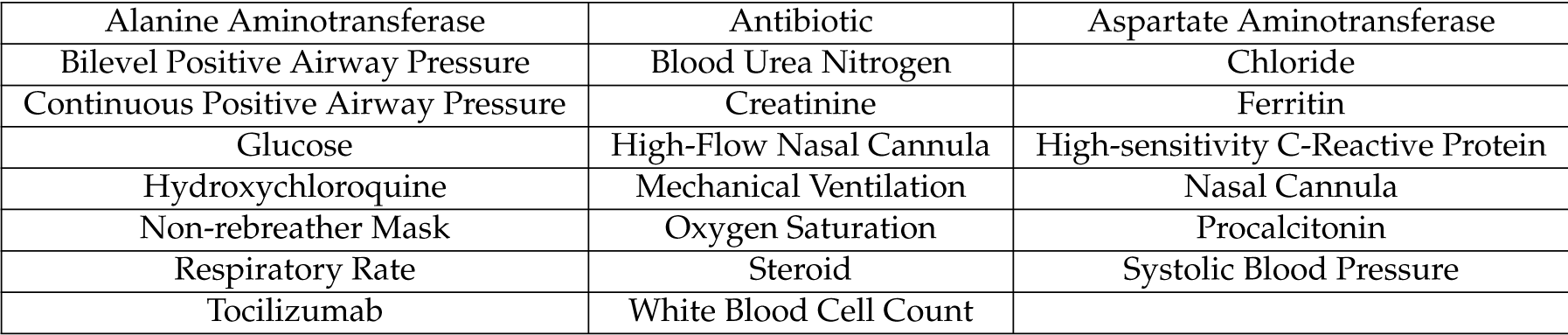
Clinical variables for the COVID-19 dataset.

|  |  |  |
| --- | --- | --- |
| Alanine Aminotransferase | Antibiotic | Aspartate Aminotransferase |
| Bilevel Positive Airway Pressure | Blood Urea Nitrogen | Chloride |
| Continuous Positive Airway Pressure | Creatinine | Ferritin |
| Glucose | High-Flow Nasal Cannula | High-sensitivity C-Reactive Protein |
| Hydroxychloroquine | Mechanical Ventilation | Nasal Cannula |
| Non-rebreather Mask | Oxygen Saturation | Procalcitonin |
| Respiratory Rate | Steroid | Systolic Blood Pressure |
| Tocilizumab | White Blood Cell Count |  |

**Table 4.** Flow cytometry markers for the COVID-19 dataset.

|  |  |  |  |  |  |  |
| --- | --- | --- | --- | --- | --- | --- |
| CCR7 | CD3 | CD4 | CD8 | CD25 | CD38 | CD45RA |
| CD127 | CXCR5 | FSC | HLA-DR | PD1 | SSC | TIM3 |

To evaluate the ability of the FMGAN to perform this generation, we train on 90% of the patients (116) and withhold 10% of the patients (13) for evaluation. We train to generate a distribution of FACS measurements from each single condition corresponding to a patient’s clinical measurements. In Figure 5, we see the resulting data from all 13 heldout patients in the top row. In the second row, we see the corresponding FMGAN generated data. Remarkably, the FMGAN learned to accurately model the true distribution of FACS data even for the never-before-seen patients. Distinct populations of cells are visible: CD3+ T cell populations including both CD4+ (T helper cells) and CD8+ (Cytotoxic T cells), as well as a CD38+ population. With each protein marker, the FMGAN accurately models the underlying data distribution.

**Figure 5.**
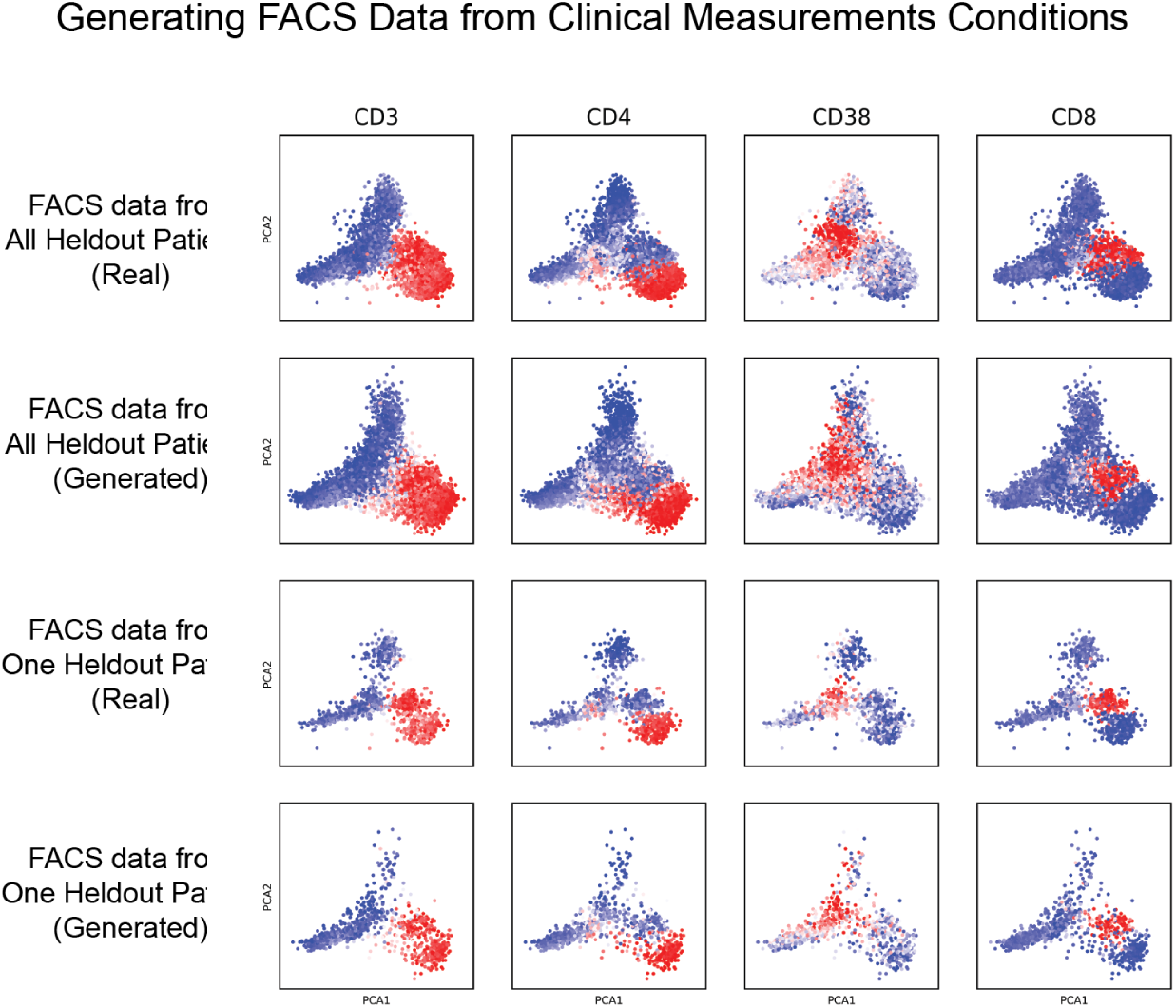
FACS data generated from clinical measurements in the COVID-19 data. Top row: for all 13 held-out patients, the real FACS measurements. Second row: for all 13 held-out patients, generated FACS measurements from the FMGAN. Third row: a single patient’s real FACS measurements. Bottom row: a single patient’s generated FACS measurements.

In the bottom two rows of Figure 5, we see the FMGAN model the distribution from a single patient accurately, as well. This per-patient generation forms the basis for our quantification of the model’s accuracy. We utilize the same baseline as in the previous section. For each patient, we measure the distribution distance between the predicted distribution and the true distribution of FACS data (scored by MMD, as before). Table 2 shows the FMGAN is able to produce distributions very close to the true underlying distribution for each patient, while the baseline model does not. As each distribution is complex with many different cell populations with varying proportions, it is not surprising that the more richly expressive FMGAN is better able to model the true data.

We note that with the FMGAN, we are able to predict the FACS measurements on never-before-seen patients, based on their clinical measurement alone. However, this relied upon the patients in the training set being representative of the patients in the held-out set. In practical applications, this means that the population of patients would need to be chosen carefully and diversely for the predictions to be meaningful for future patients.

## 3. Methods

### 3.1. Conditional Generative Adversarial Networks

In a Generative Adversarial Network (GAN), samples from the generator *G* can be obtained by taking samples from *z* ∼ *Z* and then performing the forward pass with the learned weights of the network. But while the values of *z* control which points *G* generates, we do not know how to ask for specific types of points from *G* (more discussion of the original, unconditional GAN is in the Supplementary Information).

The lack of this functionality motivated the need for the conditional GAN (cGAN) framework [18, 19]. The cGAN augments the standard GAN by introducing label information for each point. These labels stratify the total population of points into different groups. The generator is provided a given label in addition to the random noise as input, and the discriminator is provided with not only real and generated points, but also the labels for each point. As a result, the generator not only learns to generate realistic data, but it also learns to generate realistic data *for a given label*.

After training, the labels, whose meaning is known to us, can be provided to the generator to generate points of a particular type on demand. Because *G* is provided both a label and a random sample from *Z*, the cGAN is able to model not just a mapping from a label to a single point, but instead a mapping from a label to an entire distribution.

Expressing the cGAN formula mathematically yields a similar equation as to the original GAN, except with the modeled data distributions being marginal distributions conditioned on the label *l* of each point:

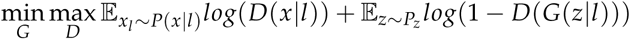

Learning a generative model conditioned on the labels allows information sharing across labels, another advantage of the cGAN framework. Since the generator *G* must share weights across labels, the signal for any particular label *l*_*i*_ is blended with the signal from all other labels *l*_*j*_, *j* = *i*, allowing for learning without massive amounts of data for each label.

### 3.2. Chemical Structure and SMILES Strings

Conditional GANs are a powerful construction for guided generation, but require some known label space to be used. While the label space must be relevant to the measured data space for an informative model to be learned, the relationship need not be simple and can be noisy. When the data space is gene expression after a drug perturbation, as in our application here, one relevant source of labels is metadata about the structure of the drug used for the perturbation. We consider two ways of representing this structure for our label space: a one-dimensional sequence of letters called a Simplified Molecular-Input Line-Entry System (SMILES) string, and a two-dimensional image called a structure diagram.

#### SMILES strings

A SMILES string encodes the chemical structure of a drug in a variable-length set of standard letters and symbols. Each character in the string represents an element of the chemical’s physical formation, for example an atom, a bond, or a ring. For example, the common molecule glucose has the following structure:

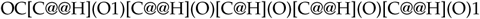

The letters indicate elements oxygen, carbon, and hydrogen, with @ denoting steriochemical configuration, and brackets and parentheses representing bonds and branches, respectively. Clearly, while providing rich information about the drug, this representation does not immediately lend itself to use as a condition. In order to distill these variable-length sequences into a fixed-size representation where similar structures have similar representations, we use a sequence-encoding neural network to embed each structure into a latent space.

#### Structure diagram

An alternate way of representing chemical structure, more intelligible for a human observer than SMILES strings, is a structure diagram. These have letters representing elements as in the SMILES strings, but also are distinguished by colors, while different types of bonds are indicated with simple lines. These images are downloaded from the PubChem PUG REST API [15]. While specifying how to get information about the structure out of this image explicitly would be impossible (in terms of RGB pixels), a neural network can learn how to process these images itself in order to accomplish its training objective, all through a completely differentiable optimization with stochastic gradient descent.

### 3.3. FMGAN

We describe the architecture for the FMGAN in this section. In the SMILES strings experiment, to obtain a fixed-length *D*_*E*_-dimensional vector for each string, we represent each input as a sequence of length *N*_*seq*_ vectors, with *N*_*seq*_ being the longest SMILES string in the database. Each element in the sequence is a vector representing the character in that position of the sequence (with a null token padding the end of any sequence shorter than *N*_*seq*_). As is standard in language processing, we learn character-level embeddings simultaneously with the sequence-level processing. Let *V* be the vocabulary, or set of all characters. The character-level embeddings are rows of a |*V*| *× D*_*char*_ matrix *W*, where |*V*| is the number of characters in the vocabulary and *D*_*char*_ is a hyperparameter, the size of the character embedding. Each input is then represented as a sequence where the *i*^*th*^ element is the row of *W* corresponding to the *i*^*th*^ character in the SMILES string.

The size of the vocabulary (number of characters including start, end, and null tokens) is 43. We chose the size of the character-level embedding to be 100. The embedder network *E* consists of two convolutional layers with 64 and 32 filters, respectively, each with a kernel-size of 40 and stride-length 2 with batch normalization and a leaky ReLU activation applied to the output. These convolutional layers are followed by four fully-connected layers which gradually reduce the dimensionality of the data with 400, 200, 100, and 50 filters, respectively. All layers except the last one have batch normalization and leaky ReLU activations. The generator and discriminator have the same architecture as the previous experiment.

This input representation is then passed through *E*, a convolutional neural network (CNN), which produces the sequence embeddings. *E* performs one-dimensional convolutions over each sequence followed by fully-connected layers, eventually outputting a single *D*_*E*_-dimensional vector for each SMILES string. We let these embeddings form the condition space for the next stage in FMGAN, the conditional GAN.

For the structure diagram experiment, we start with images that are points in *h* x *w* x *c* space, with *h* = *w* = 64 and *c* = 3. They are then processed with a CNN. The CNN consists of four convolutional layers with stride 2, kernel size 3, and filters of 32, 64, 128, and 256, respectively. Batch normalization and a ReLU activation was used for each layer. Finally, after the convolutions, one fully connected layer maps the flattened output to a 100-dimensional point, representing the embedding learned for the particular diagram.

For both experiments, the generator structure, after the drugs are processed into conditions, is the same. Let *c*_*i*_ be the condition for drug *i* formed by the embedder. Let *x*_*i*_ be the *D*_*x*_-dimensional corresponding gene expression profile from a perturbation experiment performed with drug *i*. We build a GAN that trains a generator *G* to model the underlying data distribution conditioned upon the structure *p*_*data*_(*x*|*c*). *G* takes as input both a sample from a noise distribution (we choose an isotropic Gaussian) *z* ∼ *Z*, and a condition *c*_*i*_. *G* maps these inputs to a *D*_*x*_-dimensional point. Then, the discriminator *D* takes both a *D*_*x*_-dimensional point and a condition *c* and outputs a single scalar representing whether it thinks the point was generated by *G* or was a sample from *p*_*data*_. These networks then train in the standard alternating gradient descent paradigm of GANs previously detailed.

For specific hyperparameter choices and data dimensionality details, we refer to the Supplementary Information.

We note a few additional points about the FMGAN framework. First, since everything in the network including the character-level embeddings, the embedder *E*, and the GAN are all expressed differentiably, the whole pipeline can be trained at once in an end-to-end manner. Thus, the character-level embeddings and the convolutional weights can be optimized for producing SMILES strings embeddings *useful for this specific task and context*. This is a powerful consequence, as defining what makes a good static embedding of a high-dimensional sequence may be ambiguous without reference to a particular task.

## 4. Discussion

The FMGAN model allows us to predict hard-to-obtain information for samples where we only directly measure easy-to-obtain information. We demonstrate that the FMGAN can accurately model never-before-seen samples in these contexts. In the drug discovery context, this allows the potential impact of saving on expense and time by not performing as many physical experiments and instead modeling their results. In the clinical context, this allows for the modeling of patient data sooner, with more time to take positive interventions.

Furthermore, the flexible framework of the cGAN we develop for the FMGAN allows for EI that requires advanced processing to be used as the conditional input. We demonstrate this on images and long one-dimensional sequences, but this can extended to other difficult-to-represent data. For example, in the clinical setting, the advances in natural language processing achieved by deep neural networks could be utilized to process doctor’s notes as raw text and then incorporated into the model. We demonstrate that the FMGAN is able to leverage structure in the condition space in both manifold form (from the PHATE coordinates) and discrete form (from chemical structure strings). While seemingly similar, these are very different from an information theoretical point of view. In the manifold setting, differences in input can create differences in output in a smooth way, but in the discrete setting, one small change in an individual feature may have a large effect on the output while another small change in a different feature has no effect on the output at all. For example, in a chemical structure string, modifications to some locations will not change the function at all, while in other locations a single change will determine function.

While we demonstrate that the FMGAN can be usefully applied to generative problems in a wide variety of modalities, and, as we show, even in the presence of high amounts of stochasticity.

## 5. Software Availability

https://github.com/KrishnaswamyLab/FMGAN

## 6. Supplementary Information

### 6.1. Generative Adversarial Networks

Generative Adversarial Networks (GANs) are a deep learning framework for learning a generative model of a data distribution. In recent years, they have gained significant popularity by achieving state-of-the-art performance on applications to images, language, sequences, and many other data modalities [14,20–23]. GANs differ from other types of models by not using explicit likelihood measures nor relying on having a meaningful distance measure between points. Instead, they teach a generator neural network *G* with a second discriminator network *D* using the following equation:

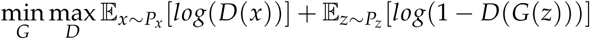

where *x* is the training data, *z* is a noise distribution that provides stochasticity to the generator and is chosen to be easy to sample from (typically an isotropic Gaussian).

### 6.2. Conditional Generative Adversarial Networks

Conditional Generative Adversarial Networks (cGANs) originated from the desire for having greater control over generation from GANs. In the case where external information, such as class labels, are available, we would like to be able to generate a random point from a specific class. The methods devised to achieve this involve providing a random label to the generator during training and then providing this label and the generated image to the discriminator. The discriminator also receives real images and their labels, allowing it to learn their joint distribution.

Once the model has been trained in this way, control over generation can be used to generate a point from a particular class by feeding the desired class into the generator. This process especially benefits from having fine-grained, continuous conditions like we have, as this gives even more precise control over generation.

### 6.3. Optimization

The networks *G* and *D* take turns optimizing their objectives through alternating gradient descent. Throughout training, the discriminator provides gradient information to the generator guiding it to better quality generation. This powerful framework provides the ability to model arbitrarily complex distributions without making any explicit parametric or limiting assumptions about their shape.

Theoretical analysis of GANs have shown their ability to converge to an optimal point where the generated distribution is indistinguishable from the true distribution [24–26]. The ability to converge to this optimal generative model without specifying a distribution distance is especially helpful in our applications, where the points lie in high dimensions and the curse of dimensionality makes distances problematic [27].

#### Manifold learning

A useful assumption in representation learning is that high biomedical dimensional data originates from an intrinsic low dimensional manifold that is mapped via nonlinear functions to observable high dimensional measurements; this is commonly referred to as the manifold assumption. In particular, we believe that since biological entities like patients, cells lie in lower dimensional spaces because of informational redundancy and coordination between measured features (coordinating genes, or coordinated combinations of residues on molecules). Further, we believe that these low dimensional spaces form smoothly varying patches because of natural heterogeneity between entities. The fact that the manifold model is successful in modeling biological entities has been shown in literature numerous times [28] and has lead to successful methods data denoising [29], clustering [30], visualization [31], and progression analysis [32].

Formally, let *M*^*d*^ be a hidden *d* dimensional manifold that is only observable via a collection of *n* ≫ *d* nonlinear functions *f*_1_, …, *f*_*n*_ : *M*^*d*^ *→* R that enable its immersion in a high dimensional ambient space as *F*(*M*^*d*^) = {**f**(*z*) = (*f*_1_(*z*), …, *f*_*n*_(*z*))^*T*^ : *z ∈ M*^*d*^} ⊆ R^*n*^ from which data is collected. Conversely, given a dataset *X* = {*x*_1_, …, *x*_*N*_} *⊂* R^*n*^ of high dimensional observations, manifold learning methods assume data points originate from a sampling 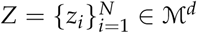 of the underlying manifold via *x*_*i*_ = **f**(*z*_*i*_), *i* = 1, …, *n*, and aim to learn a low dimensional intrinsic representation that approximates the manifold geometry of *M*^*d*^.

To learn a manifold geometry from collected data, we use the popular diffusion maps construction of [33] that uses diffusion coordinates to provide a natural global coordinate system derived from eigenfunctions of the heat kernel, or equivalently the Laplace-Beltrami operator, over manifold geometries. This construction starts by considering local similarities defined via a kernel *K*(*x, y*), *x, y ∈ F*(*M*^*d*^), that captures local neighborhoods in the data. We note that a popular choice for *K* is the Gaussian kernel exp(*-*∥*x - y*∥^2^/*σ*), where *σ >* 0 is interpreted as a user-configurable neighborhood size. However, such neighborhoods encode sampling density information together with local geometric information. To construct a diffusion geometry that is robust to sampling density variations we use an anisotropic kernel

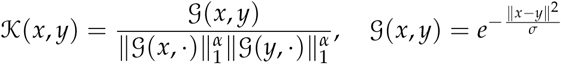

as proposed in [33], where 0 *≤ α ≤* 1 controls the separation of geometry from density, with *α* = 0 yielding the classic Gaussian kernel, and *α* = 1 completely removing density and providing a geometric equivalent to uniform sampling of the underlying manifold. Next, the similarities encoded by *K* are normalized to define transition probabilities 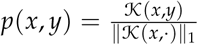 that are organized in an *N × N* row stochastic matrix

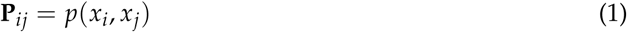

that describes a Markovian diffusion process over the intrinsic geometry of the data. Finally, a diffusion map [33] is defined by taking the eigenvalues 1 = *λ*_1_ ≥ *λ*_2_ ≥ … ≥ *λ*_*N*_ and (corresponding) eigenvectors 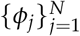 of **P**, and mapping each data point *x*_*i*_ *∈ X* to an *N* dimensional vector 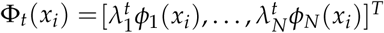, where *t* represents a diffusion-time (i.e., number of transitions considered in the diffusion process). In general, as *t* increases, most of the eigenvalues 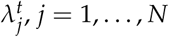, become negligible, and thus truncated diffusion map coordinates can be used for dimensionality reduction [33].

#### PHATE for structure-preserving visualization of Data

Several dimensionality reduction methods that render data into 2-D visuals like PCA and tSNE. [34] and UMAP [35] exist. However, they often cannot handle the degree of noise in biomedical data. More importantly, most of these methods are not constructed to preserve the global manifold structure of the data. PCA cannot denoise in non-linear dimensions, tSNE/UMAP effectively only constrains for near neighbor preservation—losing global structure. This motivated us to develop a method of dimensionality reduction that retains both local and global structure, and denoises data [31].

PHATE also builds upon the diffusion-based manifold learning framework described above, and involves the creation of a diffused Markov transition matrix from cellular data, as in MAGIC, **P**^*t*^ (Equation 1). PHATE collects all of the information in the diffusion operator into two dimensions such that global and local distances are retained. To achieve this, PHATE considers the *i*th row of **P** as the representation of the *ith* datapoint in terms of its *t*-step diffusion probabilities to *all* other datapoints. PHATE then preserves a novel distance between two datapoints, based on this representation called *potential distance (pdist)*. Potential distance is an *M*-divergence between the distribution in row *i*, 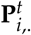 and the distribution in row *j*, 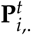. These are indeed distributions as **P**^*t*^ is 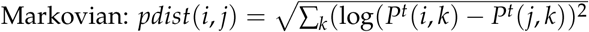

The log scaling inherent in potential distance effectively acts as a damping factor which makes faraway points similarly equal to nearby points in terms of diffusion probability. This gives PHATE the ability to maintain global context. These potential distances are embedded with metric MDS as a final step to derive a data visualization. We have shown that PHATE outperforms tSNE [34], UMAP [35], force directed layout and 12 other methods on preservation of manifold affinity, and adjusted rand index on clustered datasets, in a total of 1200 comparisons on synthetic and real datasets. In [31] we also showcased the ability of PHATE to reason about differentiation systems and differentiation trajectories in human embryonic cell development.

### 6.4. Maximum Mean Discrepancy

To evaluate the accuracy of the predicted distribution with respect to the true distribution for a given condition, we utilize maximum mean discrepancy (MMD) [36]. The MMD is a distribution distance based on a kernel applied to pairwise distances of each distribution. Specifically, MMD is calculated as:

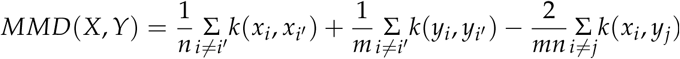

for finite samples from distributions *X* = *{x*_1_…*x*_*m*_*}* and *Y* = *{y*_1_…*y*_*n*_*}* and kernel function *k*. Two distributions have zero MMD if and only if they are equal. MMD has been used successfully in biological systems in the past, particularly in detecting whether two systems were different in distribution [37].

### 6.5. eICU Clinical data

A patient cohort at high risk for mortality due to severe illness was selected from the eICU Collaborative Research Database, a public multicenter critical care database containing 200,859 ICU admissions with 139,367 unique patients admitted to critical care units between 2014 and 2015 [8,9]. After excluding patients who did not have a calculated risk score for mortality, the APACHE IVa score and who had at least one vital sign, the final dataset contained 146,587 encounters with 118,638 unique patients. The structured datafields from automated vital signs, laboratory results, and treatments for the patient cohort were extracted and transformed as described by a recent manuscript by taking the most abnormal values in the first 24 hours from ICU admission and multiple imputation using Bayesian Ridge Regression was used to fill missing variables [38].

### 6.6. COVID-19 Clinical data

The cohort of patients included only those who were hospitalized at any of 6 hospitals in the Yale-New Haven Health System (Bridgeport, Greenwich, St. Raphael’s Campus, Westerley, Lawrence and Memorial, York Street Campus) during the period between March 1st, 2020 and June 1st, 2020 with a positive COVID test (nasopharyngeal source) between admission and discharge. Only the first encounter was included in the dataset for patients with multiple encounters during the time period of observation. Patients with a positive test prior to hospital admission but not tested during admission or tested negative during admission were not included in the cohort. Data for these patients was then extracted from the electronic health record (Epic, Verona, WI) and included data domains of demographics (e.g. age and sex), medical history (e.g. history of diabetes), laboratory samples (e.g. white blood cell count), as well as vital signs (e.g. blood pressure measurement). Pre-defined outcomes included in-hospital mortality, transfer to the intensive care unit (ICU), as well as requirement for invasive ventilation. In-hospital mortality was measured as patients being discharged from the hospital with a deceased status. ICU care was measured through location data for patients and was manually validated through chart review. Ventilation status was measured through procedure orders placed during the patient’s hospitalization and were validated through chart review.

Time-varying data, specifically vital signs as well as laboratory studies, were extracted at all timepoints of measurement during a patient’s admission.

Features were selected from a predictive model developed to predict early hospital respiratory decompensation among patients with Covid-19 and augmented with treatment received. There were a total of 19 clinical, laboratory, and treatment variables extracted: systolic blood pressure, respiratory rate, oxygen saturation, blood urea nitrogen, creatinine,chloride, glucose, white blood cell count, alanine aminotransferase, aspartate aminotransferase, high-sensitivity C-reactive protein, ferritin, procalcitonin, age, gender, and treatment with hydroxychloroquine, steroid, antibiotic, or tocilizumab. Only complete cases, or patients with recorded values for all 19 variables in the first 24 hours, were included in the final dataset.

As preprocessing, the most abnormal value in the first 24 hours was selected for the clinical and laboratory variables according to the methodology described in a previous electronic health record-based study. The categorical variables for treatment were coded as binary (1 for received, 0 for not recorded).

## Notes

### Competing Interest Statement

The authors have declared no competing interest.

